# EBV miRNAs are potent effectors of tumor cell transcriptome remodeling in promoting immune escape

**DOI:** 10.1101/2020.12.21.423766

**Authors:** Nathan Ungerleider, Whitney Bullard, Mehmet Kara, Xia Wang, Claire Roberts, Rolf Renne, Scott Tibbetts, Erik Flemington

**Affiliations:** Department of Pathology, Tulane University School of Medicine, Tulane Cancer Center, New Orleans, LA 70112, United States of America; Dept. of Molecular Genetics and Microbiology, UF Health Cancer Center, University of Florida, Gainesville, Florida, United States of America

**Author notes:** To whom correspondence should be addressed. Erik Flemington, Correspondence may also be addressed to: Scott Tibbetts.

## Abstract

Epstein Barr virus (EBV) contributes to the tumor phenotype through a limited set of primarily non-coding viral RNAs, including 31 mature miRNAs. Here we investigated the impact of EBV miRNAs on remodeling the tumor cell transcriptome. Strikingly, EBV miRNAs displayed exceptionally abundant expression in primary EBV-associated Burkitt’s Lymphomas (BLs) and Gastric Carcinomas (GCs). To investigate viral miRNA targeting, we used the high-resolution approach, CLASH in GC and a BL cell models. Affinity constant calculations of targeting efficacies for CLASH hits showed that viral miRNAs bind their targets more effectively than their host counterparts, as did Kaposi’s sarcoma-associated herpesvirus (KSHV) and murine gammaherpesvirus 68 (MHV68) miRNAs. Using public BL and GC RNA-seq datasets, we found that high EBV miRNA targeting efficacies translates to enhanced reduction of target expression. Pathway analysis of high efficacy EBV miRNA targets showed enrichment for innate and adaptive immune responses. Inhibition of the immune response by EBV miRNAs was functionally validated *in vivo* through the finding of inverse correlations between EBV miRNAs and immune cell infiltration and T-cell diversity in TCGA GC dataset. Together, this study demonstrates that EBV miRNAs are potent effectors of the tumor transcriptome that play a role in suppressing the host immune response.

**AUTHOR SUMMARY:** Burkitt’s Lymphoma and gastric cancer are both associated with EBV, a prolific DNA tumor virus that latently resides in nearly all human beings. Despite mostly restricting viral gene expression to noncoding RNAs, EBV has important influences on the fitness of infected tumor cells. Here, we show that the miRNA class of viral noncoding RNAs are a major viral contributor to remodeling the tumor cell regulatory machinery in patient tumor samples. First, an assessment of miRNA expression in clinical tumor samples showed that EBV miRNAs are expressed at unexpectedly high levels relative to cell miRNAs. Using a highly specific miRNA target identification approach, CLASH, we comprehensively identified both viral and cellular microRNA targets and the relative abundance of each microRNA-mRNA interaction. We also show that viral microRNAs bind to and alter the expression of their mRNA targets more effectively than their cellular microRNA counterparts. Pathway analysis of the most effectively targeted mRNAs revealed enrichment of immune signaling pathways and we show a corresponding inverse correlation between EBV miRNA expression and infiltrating immune cells in EBV positive primary tumors. Altogether, this study shows that EBV miRNAs are key regulators of the tumor cell phenotype and the immune cell microenvironment.

## INTRODUCTION

The Epstein Barr Virus (EBV) is a ubiquitous gammaherpesvirus (*γ*HV) that establishes lifelong infections in over 90% of the world’s population. Since its discovery as the major etiological agent of endemic Burkitt’s lymphoma (BL)(1), EBV has been causally associated with other malignancies including NK/T cell lymphoma(2), diffuse large B cell lymphoma(3), Hodgkin’s lymphoma(4), nasopharyngeal carcinoma(5) and gastric carcinoma (GC)(6). Arising from infected founder cells, these tumors maintain a dependence on the virus as they progress(7).

*γ*HVs utilize two distinct strategies, referred to as lytic and latent replication, to expand the infected host cell population(8). The lytic replication program is a process utilized by all viruses to replicate and package their genetic content for spread through cell-to-cell transfer. This strategy produces a large number of virions but it destroys the host cell and triggers a strong local immune response(9). The viral latency program, the phase most closely linked with cancer, is associated with the expression of a small number of genes that support the growth and health of the infected cell(10-16) and by extension, the growth and health of its viral occupant. In this phase of the virus infection cycle, viral and host genomes are replicated concordantly and distributed to daughter cells(17), resulting in an expansion of the infected cell population that is independent of virion production. This form of intracellular replication minimizes immune reactivity, allowing the virus to discretely double the pool of infected cells at each mitotic cycle.

In EBV-associated BLs and GCs, the latency gene expression program is especially restrictive, with the only ubiquitous and consistently expressed viral protein being the replicative factor, EBNA1(18). More abundantly expressed are a group of viral noncoding RNAs that allow the virus to modulate the host cell environment without eliciting a strong adaptive immune response. These include two short noncoding RNAs (EBER1 and EBER2)(19), a long noncoding RNA (RPMS1)(20), circular RNAs(21, 22), and the viral miRNAs(23).

MiRNAs function by targeting complementary mRNAs for destruction(24). By interacting with an average of 90 unique transcripts(25), a single miRNA can have a substantial impact on the cellular transcriptome(26-30). The genomes of EBV, KSHV, and MHV68 potentially encode 44, 25, and 28 miRNAs(31-34), theoretically endowing these viruses with the capacity to control nearly every pathway in the cell. Several previous studies have used broad-scale approaches such as AGO-CLIP to uncover viral miRNA targets(35-37), and some of these miRNA-target interactions have since been shown to interrupt apoptosis(38), prevent reactivation(39), and block interferon signaling(40). The AGO-CLIP approach, however, requires bioinformatic inferences to determine miRNA-mRNA pairings, precluding analysis of binding efficacy and limiting detection to canonical interactions. Here we used a modified version of a more comprehensive and quantitative approach, Crosslinking, Ligation, and Sequencing of Hybrids (CLASH)(41), called qCLASH(42), to broadly uncover *bona fide* targets of EBV miRNAs. Through integration of mRNA, miRNA, and hybrid abundances, we examine the global binding properties and targeting efficacies inherent to *γ*HV miRNAs. Assessing pathway enrichment of the highest targeting efficacy EBV miRNA-target interactions shows enrichment for innate and adaptive immune responses. Utilizing clinical BL and GC datasets, we explore the roles of EBV miRNAs in primary tumors and demonstrate a role for EBV miRNAs in dampening the immune response to viral infection and oncogenesis.

## MATERIALS AND METHODS

### Cell culture

Akata (obtained from Kenzo Takada) and SNU719 (Korean Cell Line Bank) cells were grown in RPMI 1640 media (ThermoFisher Scientific, catalog no. SH30027) supplemented with 10% fetal bovine serum (FBS; ThermoFisher Scientific, catalog no. 10437), and incubated at 37°C in 5% CO_2_.

### Clinical data

Aligned RNA and miRNA sequencing reads, deposited by the BGLSP and TCGA, were downloaded from the Genomic Data Commons (GDC)(43). RNA sequencing alignments were converted to FASTQ format using the following command:

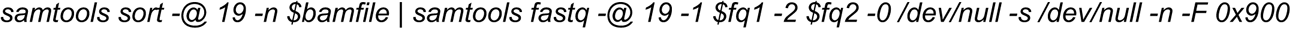

The resulting raw sequencing reads were pseudoaligned to the human (GENCODE GRCh38.p13)(44) and EBV(45) transcriptomes using kallisto v0.46.0(46) (including the “-rf-stranded” flag for BL sequencing reads, which were strand-specific). The underlying counts or transcripts per million (t.p.m.) values were summed and assigned to each gene. BL analysis was restricted to the “Discovery” cohort due to the reported higher RNA sequencing quality of these samples(47). With the exception of *Figure S1*, GC analysis was restricted to microsatellite stable samples(48).

Aligned miRNA sequencing reads were converted to FASTQ format using the following command:

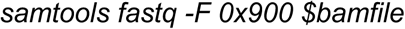

Raw miRNA sequencing reads were aligned to an index of mature human and EBV miRNA sequences (miRbase v22) via bowtie with the following command:

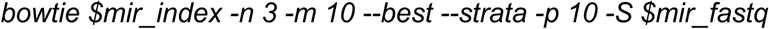

### UV Crosslinking

qCLASH was performed as previously described^1^, with 3 biological replicates processed for each cell line. SNU719 cells were trypsinized, resuspended in RPMI 1640 plus 10% FBS, then washed twice with PBS. Akata cells were washed twice with PBS. Both cell lines were resuspended in 10 mL PBS then crosslinked with 250 nm λ UV light for a total exposure of 600J/cm^2^. 50 million cells were transferred to a 1.5 mL tube, pelleted at 800 × g for 5 minutes, and stored at −80°C before processing as described below.

### Cell lysis

Cell pellets were thawed and resuspended in 500 µl cell lysis buffer (50 mM HEPES (pH 7.5), 150 mM KCl, 2mM EDTA, 1mM NaF, 0.5% NP-40, 0.5 mM DTT, and 1x protease inhibitor cocktail (Roche, catalog no. 11836170001)). Cells were lysed for 15 minutes on ice then treated with 10 µL of RQ1 DNase (Promega, catalog no. M610A) for 5 minutes at 37°C, while shaking at 1,000 rpm. Lysates were cleared by a 21,000 × g centrifugation for 15 minutes at 4°C. Supernatants were treated for 15 minutes with 0.5 µL RNase T1 (ThermoFisher Scientific, catalog no. EN054) at 22°C.

### Preparation of Protein G beads

6 mg of Protein G Dynabeads (Invitrogen, catalog no. 1004D) were washed 3 times then resuspended in PBST (pH 7.2). A final concentration of 200ug/ml of AffiniPure rabbit α-mouse IgG (Jackson ImmunoResearch, catalog no. 315-005-008) was added and beads were incubated on a rotator for 50 minutes at 25°C. The complexes were washed 3 times with PBST (5 minutes each), then resuspended in 1 mL of PBST. 10 µg of pan-AGO antibody (gift from Zissimos Mourelatos) was added and samples were rotated for 16 hours at 4°C. Beads were then washed 4 times with wash buffer (1x PBS, 0.1% SDS, 0.5% sodium deoxycholate, and 0.5% NP-40).

### Immunoprecipitation

1 mL of crosslinked cell lysate was added to the pelleted α-AGO beads. Tubes were put on a rotator and incubated for 16 hours at 4°C. The bead complexes were then centrifuged and washed 3 times with cell lysis buffer, followed by 4 times each with 1x PXL (1x PBS, 0.1% SDS, 0.5% sodium deoxycholate, 0.5% NP-40), 5x PXL (5x PBS, 0.1% SDS, 0.5% sodium deoxycholate, 0.5% NP-40), high-stringency wash buffer (15 mM Tris-HCl (pH 7.5), 5 mM EDTA, 2.5 mM EGTA, 1% Triton X-100, 1% sodium deoxycholate, 0.1% SDS, 120 mM NaCl, 25 mM KCl), high salt wash buffer (15 mM Tris-HCl (pH 7.5), 5 mM EDTA, 2.5 mM EGTA, 1% Triton X-100, 1% sodium deoxycholate, 0.1% SDS, 1 M NaCl), then polynucleotide kinase (PNK) buffer (50 mM Tris-HCl (pH 7.5), 10 mM MgCl2, 0.5% NP-40).

### Ligation of RNA ends

RNA 5’-ends were phosphorylated using 4 µl T4 PNK (NEB, catalog no. M0201) in a solution consisting of 8 µl 10x PNK buffer, 2 µl RNasin Plus (Promega, catalog no. N2615), 0.8 µl 100 mM ATP (ThermoFisher Scientific, catalog no. R0041), and 65.2 µl ddH_2_O. Bead complexes were incubated for 40 minutes at 10°C then washed 3 times in PNK buffer. RNAs were then ligated by rotating for 16 hours at 4°C in 500 µl of RNA ligation solution (50 µl 10x T4 RNA ligase buffer, 60 µl 50% polyethylene glycol 8000, 125 µl 4M KCl, 12.5 µl RNasin Plus, 5 µl 100 mM ATP, 321.25 µl ddH_2_0, and 50 µl T4 RNA ligase 1 (NEB, catalog no. M0204). Bead complexes were then washed 3 times with 1 mL PNK buffer. Next, 80 µl of dephosphorylation solution (8 μl 10x dephosphorylation buffer, 2 μl RNasin Plus, 67 μl ddH2O, and 3 μl alkaline phosphatase (Roche, catalog no. 10713023001)) was added to the beads, and RNA dephosphorylation was achieved by incubating samples for 40 minutes at 10°C, with intermittent 1,000 rpm shaking every 2 minutes for 15 seconds. The bead complexes were washed 2 times with 1 mL EGTA buffer (50 mM Tris-HCl (pH 7.5), 20 mM EGTA, 0.5% NP-40), and then three times with 1 mL PNK buffer.

### Ligation of 3’-adapter

To ligate the miRCat-33 3’-linker (5′-TGGAATTCTCGGGTGCCAAGG-3′) to the newly formed RNA hybrids, beads were incubated with 42 μl ddH2O, 8 μl 10× T4 RNA ligase buffer, 16 μl 50% PEG-8000, 2 μl RNasin Plus, 8 pM of linker, and 4 μl T4 RNA ligase 2 (NEB, catalog no. M0239) for 16 hours at 16°C, with 1,000 rpm shaking every 2 minutes for 15 seconds. Bead complexes were then washed 3 times in PNK buffer.

### Elution and RNA extraction

Bead complexes were incubated in 100 µl of elution buffer (100 mM NaHCO3, 1% SDS) for 15 minutes at 25°C on a 1,400 rpm shaker. After spinning at 1,000g for 1 minute, the elution buffer was transferred to a new tube. An additional 100 µl of elution buffer was added to the bead complexes, elution was repeated and the eluates were then combined. To improve RNA phase separation of crosslinked RNA-AGO complexes, proteins were digested using proteinase K. 10 µl of proteinase K (Roche, catalog no. 03115887001) in 40 µl of proteinase K buffer (100 mM Tris-HCl (pH 7.5), 50 mM NaCl, 10 mM EDTA) was added to the 200 µl of eluate and samples were incubated at 37°C for 20 min. Samples were then subjected to phenol-chloroform extraction to purify the RNA.

### CLASH library preparation

RNA ends were phosphorylated, followed by ligation of the Solexa 5’ linker (invddT-GTTCArGrAr-GrUrUrCrUrArCrArGrUrCrCrGrArCrGrArUrC-OH). RNA was re-extracted via phenol-chloroform, as described above. Sequencing libraries were prepared using the Illumina TruSeq Small RNA Sample Prep kit according to manufacturer’s instructions, and cDNA was generated using SuperScript III reverse transcriptase (Invitrogen, catalog no. 18080093). The resulting samples were sequenced on an Illumina HiSeq 2500 machine.

### RNA sequencing

RNA was extracted from Akata and SNU719 cells using TRIzol (ThermoFisher Scientific, catalog no. 15596026). Extraction was performed according to the manufacturer’s instructions with one additional step included to improve the purity of the RNA: following isopropanol precipitation, RNA was reconstituted in 200 µl ddH_2_O with 20 µl 3M sodium acetate and 500 µl ethanol, stored at −20°C for 16 hours, spun down at 20,000 × g for 30 minutes at 4°C, and then resuming the manufacturers protocol with the 70% ethanol wash step. Small fraction sequencing libraries were prepared using the Illumina TruSeq Small RNA Library Prep Kit (Illumina, catalog no. RS-200-0012), and poly-A sequencing libraries (HE2.1 and TIVE cells) were generated using the TruSeq RNA Sample Prep Kit (Illumina, catalog no. FC-122-1001).

### Bioinformatic analysis

Adapter sequences were removed from raw sequencing reads using Trimmomatic^2^. A transcriptome index was generated for *bowtie2*^3^ alignment, as previously described^4^, with the following modifications: a. GRCh38 transcript sequences were used (obtained from Gencode^5^; version 33), b. Annotated viral transcripts^6^ and miRNA sequences (obtained from mirBase^7^) were included in the index. Processed reads were aligned to the human and viral transcriptome indexes using *bowtie2* v2.3.3; alignments run with the following parameters: -D 20 -R 3 -N 0 -L 16 -k 20 --local -i S,1,0.50 --score-min L,18,0 --ma 1 --np 0 --mp 2,2 --rdg 5,1 --rfg 5,1). Hybrid alignments were retrieved from aligned reads using the *hyb* pipeline^8^ (https://github.com/gkudla/hyb). The structure and binding energies of each hybrid was predicted using the *RNAcofold* algorithm of the *Vienna RNA* package^9^.

### Targeting efficacy

Targeting efficacy for each miRNA-mRNA hybrid was calculated using the following equation:

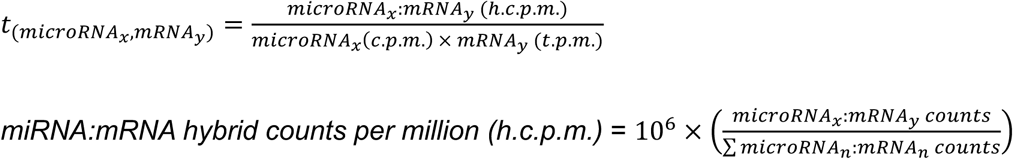

Hybrid counts were obtained via CLASH, mRNA t.p.m. were obtained from RNA-sequencing, and *miRNA c*.*p*.*m*. levels were quantified via small fraction sequencing.

### Principal component analysis

BL gene expression counts were transformed using the *normTransform* function of the *DESeq2* R package(49). GC gene expression counts were transformed using the *vsd* function of the *vsn* R package(50). Transformed counts were ranked by variance, and PCA was applied to the top 500 most variable genes in each tumor type using the *plotPCA* function of *DESeq2*. Each of the resulting principal components were compared to each relevant clinical covariate via logistic regression. *Pseudo-R*^*2*^ values(51) were calculated for each principal component and covariate pair using the *PseudoR2* of the *desctools* R package.

### Pathway analysis

CLASH derived mRNA targets for each virus and corresponding host were ranked by targeting efficacy in each cell line. Protein-protein interaction (PPI) and expression correlation networks were assembled for each of the top 20 targets using STRING(52) with default parameters, limited to 20 direct and 10 additional node proteins per queried “seed” gene. The resulting genes were explored for pathway enrichment with the *enrichR*(53) API, interrogating the Reactome library of pathways. Pathway enrichment was considered significant if the adjusted P-value < 0.05. All significant pathways were assigned to each seed gene.

## RESULTS

### High expression of viral miRNAs in EBV-associated tumors

BLs and GCs are aggressive tumors with distinct etiologies. Pediatric BL is a B-cell malignancy endemic to sub-Saharan Africa(47) that is characterized by the t(8;14)(q24;q32) *MYC:IGH* translocation(54, 55). GCs encompass a diverse group of epithelial tumors(56) originating from the stomach lining that have a broader epidemiology(57). Despite overt differences in pathology and etiology, nearly all endemic BLs (over 90%)(47) and a subset of GCs (∼10%)(6) are causally infected with EBV. As a first assessment of the viral contributions to these tumors, we performed principle component analysis (PCA) of 86 BL(47) (66 EBV+) and 235 GC(58) (24 EBV+) cell transcriptomes. Applying logistic regression to each of the top 10 (BL) or 15 (GC) principal components revealed that EBV status is a major distinguishing clinical covariate in both tumor types (*Figure 1A; Figure S1*), indicating that EBV is likely a key determinant in shaping the tumor transcriptome.

**Figure 1.**
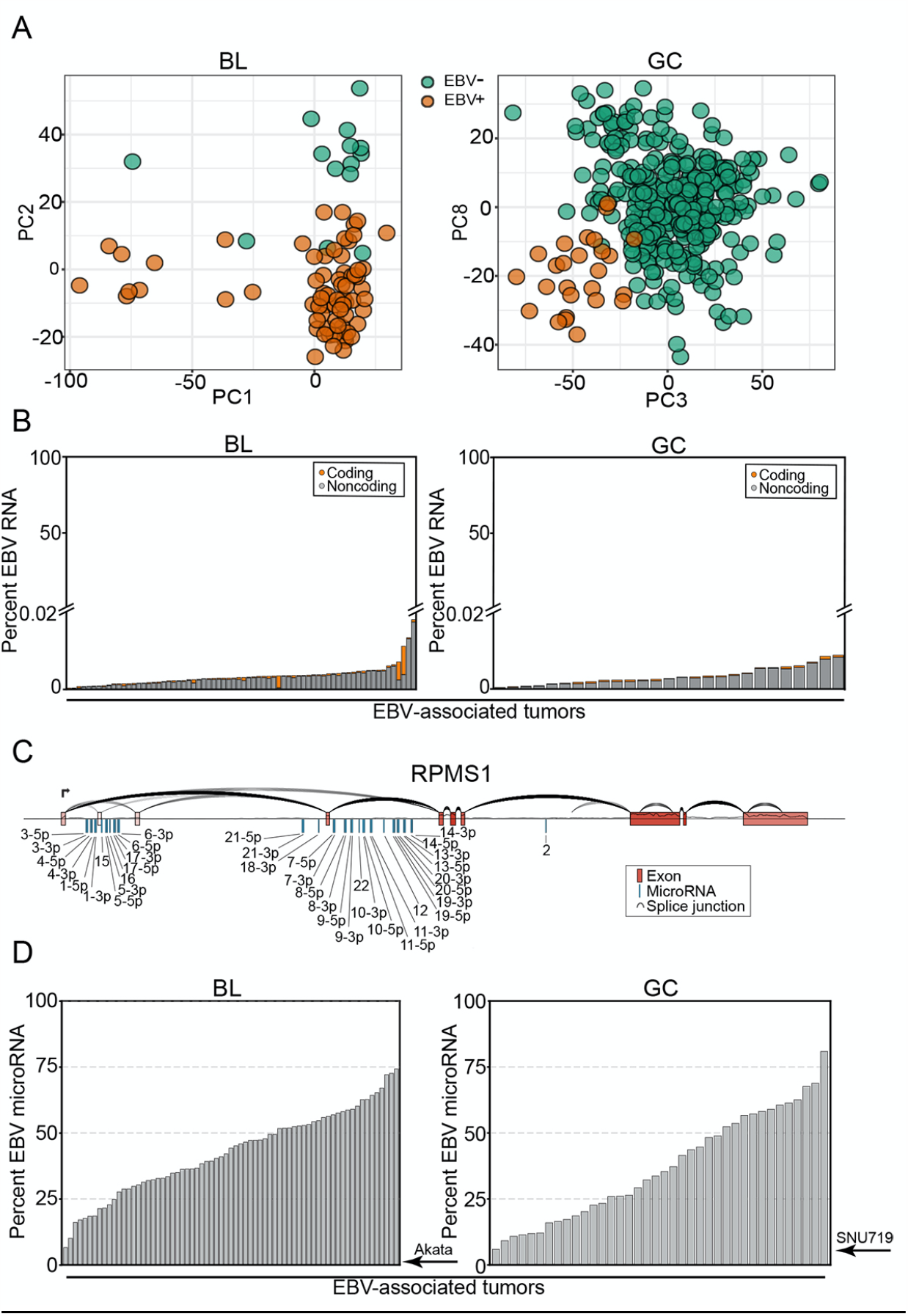
High expression of viral miRNAs in EBV-associated tumors. Raw RNA sequencing reads were obtained from the Burkitt Lymphoma Genome Sequencing Project(106) (BL; N=84) and The Cancer Genome Atlas(58) (GC; N=235). Samples were aligned to the combined human (GENCODE GRCh38.p13)(44) and EBV(45) transcriptomes. (A) Principle component analysis (PCA) of BL and GC gene expression. (B) The viral percentage of total gene expression in each tumor sample,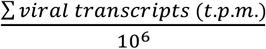 The structure of the EBV RPMS1 locus, including the BART miRNA clusters, plotted with SpliceV(107). The coverage track and splice junction counts were derived from the aligned RNA-Seq reads of an EBV^+^ BL patient (BLGSP-71-06-00281). (D) The viral percentage of total miRNA expression in each tumor sample,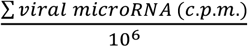. Akata and SNU719 labels indicate the viral percentage of reads in Akata and SNU719 cell lines.

To investigate underlying features of EBV that contribute to the tumor phenotype, we first carried out a quantitative assessment of the viral transcriptomes in EBV-associated BLs and GCs to identify candidate viral effectors in the natural tumor setting. Unlike lytic conditions, where 20% of expressed poly-ad-enylated RNAs in the cell are of viral origin (*Figure S2*), the sum of expressed latency transcripts in BLs and GCs does not exceed 0.02% of cellular RNAs (*Figure 1B, Figure S3*), with the majority of these being non-coding. Further, some of the EBV transcript signal in this analysis is derived from lytic gene expression occurring in a minor population of tumor cells through occasional sporadic reactivation. Therefore, long RNA latency viral gene expression represents only a minor portion of the combined cell and viral transcriptome. We were unable to assess expression of the highly abundant non-polyadenylated small noncoding viral EBER1 and EBER 2 transcripts because the GC dataset was polyA-selected and because they are partially lost following RNA size selection in the ribodepletion-derived BL dataset (which results in unreliable quantifications across samples).

Housed within the most abundantly expressed viral polyadenylated RNA, the long non-coding RNA (lncRNA), RPMS1, are 20 densely clustered intronic pre-miRNAs encoding at least 31 mature miRNAs (*Figure 1C, Figure S4-5*). In contrast to the limited contribution of viral long RNAs to the tumor transcriptomes, the cumulative expression of viral miRNAs is remarkably high, exceeding host totals by as much as 3-fold (*Figure 1D*). This is particularly relevant because miRNA function is critically dependent on sufficient miRNA expression to saturate a high fraction of each target transcript. The importance of miRNA expression levels on function is supported by previous studies that showed that poorly expressed miRNAs have little discernable biological activity(59) whereas sufficient expression of an individual miRNA can have a marked impact on tumor biology(60). The overwhelming expression of 31 viral miRNAs (*Figure 1D, Table S1*) in EBV-positive BLs and GCs supports their likely relevance in modulating the tumor transcriptome landscape.

### EBV miRNAs are over-represented in mRNA-bound RNA-induced silencing complexes (RISCs)

With the dependence on miRNA loading into RISC for target destabilization, the level of miRNA-RISC association is a predictor of miRNA targeting(61). To assess RISC association characteristics of EBV and host miRNAs, we performed a modified version(42) of Crosslinking, Ligation, And Sequencing of Hybrids (CLASH)(62), referred to as qCLASH(42), in EBV+ cell lines modeling BL (Akata) and GC (SNU719; *Figure S6A*). In CLASH, each AGO-bound miRNA-mRNA pair is ligated and then sequenced as a contiguous read, reproducibly resolving both the diversity of miRNA targets as well as the relative abundance of each interaction (*Figure S6B*).

In conjunction with our qCLASH analyses, we also performed standard small RNA fraction sequencing to assess the underlying expression of each miRNA in these cell lines. Notably, viral miRNA expression in Akata and SNU719 cell lines was substantially lower than the levels observed in primary tumors (*Figure 2A-B, left*). Nevertheless, this finding is in-line with previous studies showing that EBV miRNA levels are low in cell lines (including SNU719 cells) but increase markedly upon passage in immunocompromised mice(63, 64), likely due to an enhanced reliance on viral miRNAs in the *in vivo* setting. Despite the modest expression of viral miRNAs in *in vitro* cultured SNU719 and Akata cells, we found a remarkably high representation of EBV miRNA-mRNA hybrids (*Figure 2A-B, left)*. This observed enrichment was not attributable to a limited number of individual viral miRNAs with unusually high binding efficiencies, but instead was a characteristic that was broadly attributable to the bulk of EBV miRNAs (*Figure 2A-B, right*). These results indicate that in addition to exhibiting high *in vivo* expression, EBV miRNAs evolved with feature(s) that enhance RISC formation, supporting the possibility of distinctly productive viral miRNA targeting.

**Figure 2.**
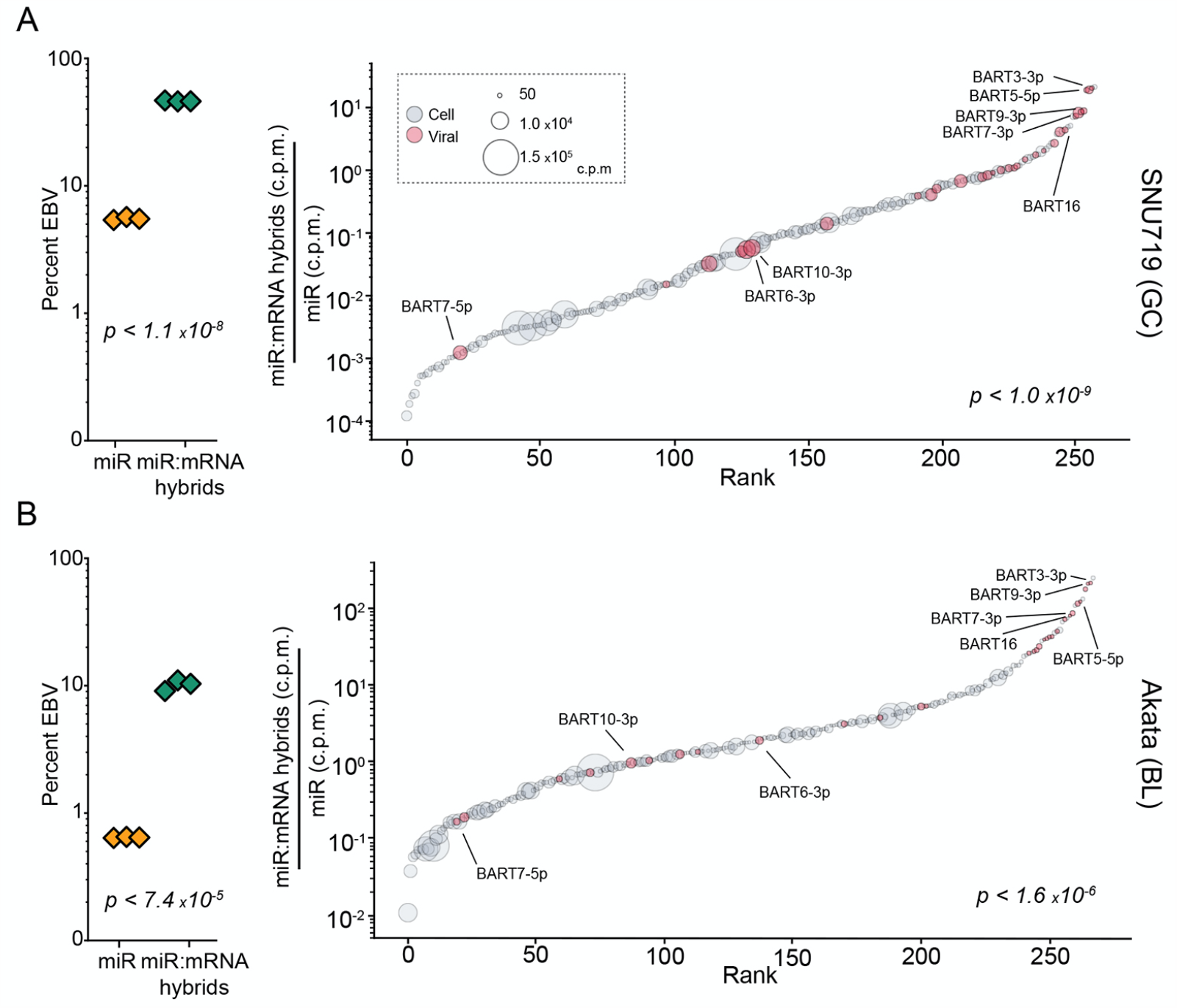
EBV miRNAs are over-represented in RNA induced silencing complexes (RISCs). CLASH and miRNA-sequencing were performed in triplicate for SNU719 and Akata cells. (A-B, left) The viral percentage of total miRNA expression in each sample, 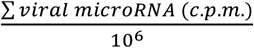 (yellow triangles), and the percent of all miRNA-mRNA hybrids containing a viral miRNA, 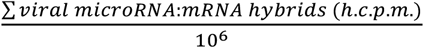 (green triangles), were calculated. The indicated P-values were calculated using unpaired Student’s t-tests. (A-B, right) The average number of miRNA-mRNA hybrids formed for each miRNA was normalized to its baseline expression level, 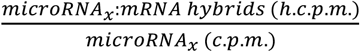. These mRNA-bound proportions were plotted in order of rank on the corresponding x-axis. Each circle represents an individual miRNA; circle size represents expression level; red circles indicate viral miRNAs. The indicated P-values were calculated using the Kolmogorov–Smirnov test (KS), comparing viral and human miRNAs.

To determine whether enhanced representation of viral miRNAs in RISC is a conserved feature of *γ*HVs, we supplemented data from our previously published KSHV(42) and MHV68(65) qCLASH studies with small RNA sequencing to similarly compare KSHV and MHV68 hybrid levels to their respective miRNA expression. Similar to our findings with *in vitro* EBV miRNA expression, KSHV miRNAs were found to be expressed at low levels in KSHV infected LTC-TIVE cells (*Figure S7A*). Like our observations with EBV miRNAs, KSHV miRNAs exhibited marked enrichment in RISCs that was distributed across a spectrum of KSHV miRNAs (*Figure S7A*). In contrast, an aggregate assessment of MHV68 miRNAs did not show over-representation within RISCs (*Figure S7B*). This lack of aggregate MHV68 miRNA enrichment in RISC, however, may be related to their non-canonical processing mechanisms(66). Nevertheless, at the individual miRNA level, we found that the two most overrepresented miRNAs within RISC in the MHV68-infected cell line, HE2.1, were the MHV68 M1-2-3p and M1-6-5p miRNAs (*Figure S7B, right*), suggesting that at least some MHV68 miRNAs are overrepresented in RISC. Together, these findings show preferential loading of viral miRNAs into RISC across these three *γ*HVs, representing a possible common strategy to exert strong influences on infected cell transcriptomes.

### EBV miRNAs have high targeting efficacies

The disproportionately high association of viral miRNAs with RISCs could result from intrinsic properties of viral miRNAs that enhance loading into RISC. Nevertheless, high expression of viral miRNA and/or their targets could also contribute to the detection of greater numbers of miRNA-target RISCs. To account for the impact of miRNA and mRNA abundance on the level of AGO-bound miRNA-mRNA hybrids(67), we used an affinity constant calculation (dividing normalized hybrid counts by the normalized number of miR-NAs and mRNA target transcripts *Figure 3C*)) as a quantitative metric for miRNA “targeting efficacy”. We first assessed how accurately this targeting efficacy metric reflects intrinsic properties of miRNAs rather than environmental, cell specific factors. Unlike cellular miRNAs, nearly all viral miRNA targets identified in Akata cells were also detected in SNU719 cells (*Figure 3A*), with the higher overall viral miRNA expression in SNU719 accounting for the additional miRNA targets identified in this system. Despite the overlap of specific targets, the relative abundance of each viral miRNA-mRNA hybrid detected in both cell lines was poorly correlated (*Figure 3B*), likely due to the differences in mRNA and/or miRNA expression in SNU719 and Akata cells. In contrast, viral miRNA targeting efficacies were strongly correlated between cell lines (*Figure 3D*). This finding also extended to cell miRNAs using miRNA/target interactions found in both cell lines (*Figure 3D*). These data support our contention that this targeting efficacy metric is a quantitative measure of intrinsic properties of miRNAs and their target sites that influence RISC formation.

**Figure 3.**
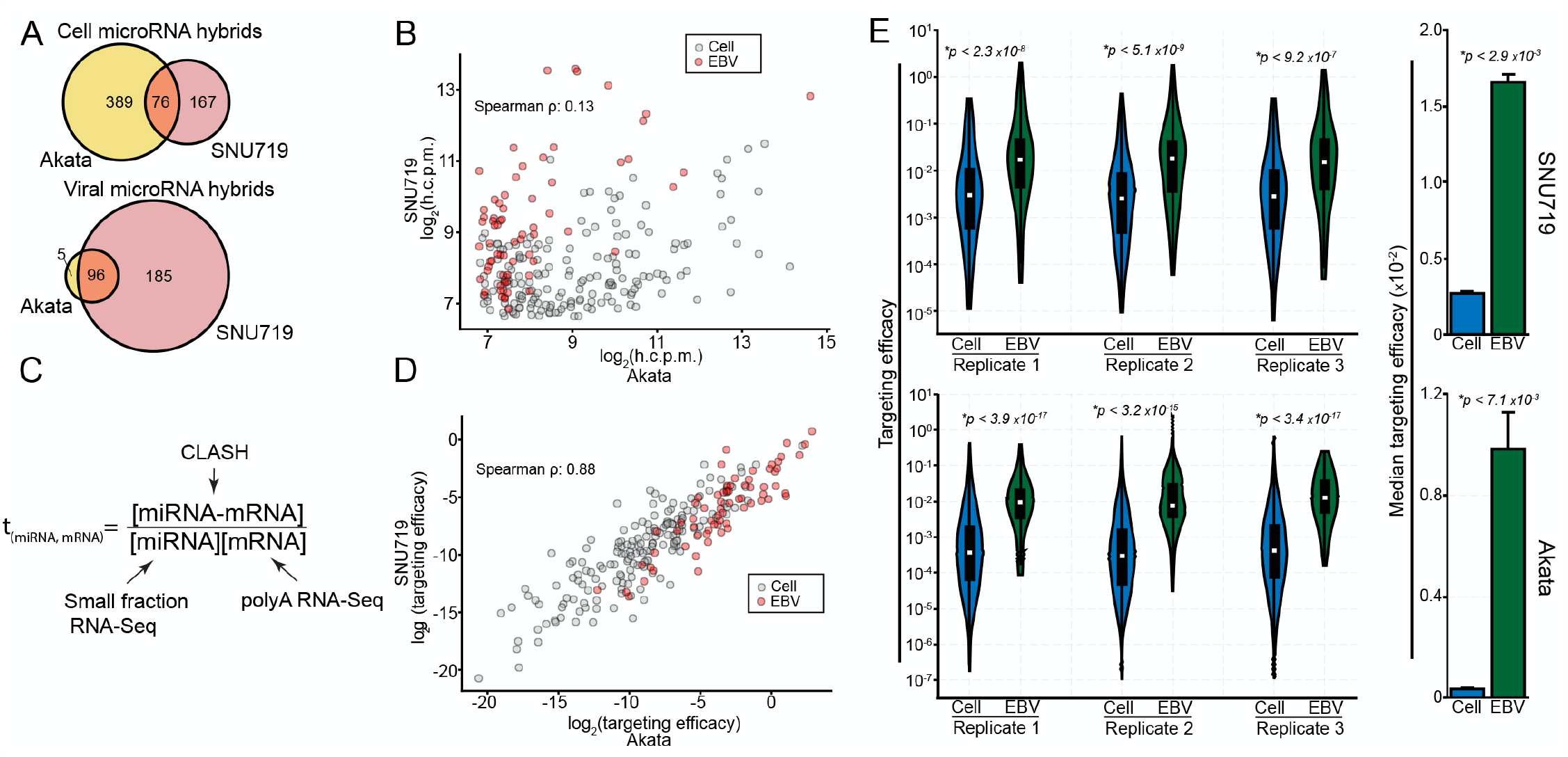
EBV miRNAs have high targeting efficacy. (A) Venn diagrams comparing genes found to be targeted in Akata and SNU719 cells by cellular (top) and viral (bottom) miRNAs. Only high confidence targets (on average >100 h.c.p.m.) were considered. (B) h.c.p.m values for each interaction pair were compared between Akata and SNU719 cells, considering all interactions found in both cell lines. The log_2_-transformed values from each cell line were correlated, resulting in a correlation coefficient of ρ = 0.13 (Spearman). (C) Schematic of the targeting efficacy calculation. (D) The log_2_-transformed targeting efficacies were correlated between SNU719 and Akata cells, resulting in a spearman correlation coefficient of ρ = 0.81. (E; left) The distribution of targeting efficacies for each interaction in each CLASH replicate, comparing viral and cellular interactions. P-values were calculated using the KS test. (E; right) The median targeting efficacy of all three replicates. P-values were calculated using paired Student’s t-tests.

We next applied targeting efficacy measures to assess the innate RISC loading properties of viral and cellular miRNAs in Akata and SNU719 cells. Across each biological replicate, the targeting efficacy distributions were substantially higher for EBV miRNAs than cell miRNAs in both SNU719 and Akata cells (*Figure 3E, Table S2*). Extending this analysis to KSHV and MHV68 miRNAs, we similarly found that viral miRNAs have higher targeting efficacies than their cellular counterparts (*Figure S8A-B, Table S2*). These results indicate that the higher proportion of viral miRNAs in RISCs relative to their expression levels is due to intrinsic properties of viral miRNAs that influence target selection and RISC loading. This suggests that viral miRNAs have evolved to be more effective inhibitors of their mRNA targets than host miRNAs.

### EBV miRNAs target more accessible regions of mRNAs than cell miRNAs and form more thermo-dynamically stable hybrids

To assess biophysical properties underlying efficient viral miRNA RISC formation, we explored the nucleotide composition of miRNA/mRNA hybrids. Hybrid formation is guided in large part by the degree of complementarity between bases 2-8 of miRNAs and their targets, with loading into RISC being further boosted by the presence of an “A” opposite the first base of the miRNA(68). These bases, collectively referred to as the “seed” region, display distinct complementarity patterns that are categorized into 9 different classes. With seed class being a known determinant of targeting effectiveness, we first tested whether there is a seed class bias for EBV miRNAs relative to cell miRNAs. As shown in *Figure S9*, there is no discernable enrichment of EBV miRNAs in seed classes with higher known targeting capabilities. Further, we found that within each seed class, viral miRNAs have higher targeting efficacies (*Figure 4A*). These data indicate that differential seed class utilization does not explain the increased targeting efficacy of EBV miRNAs.

**Figure 4.**
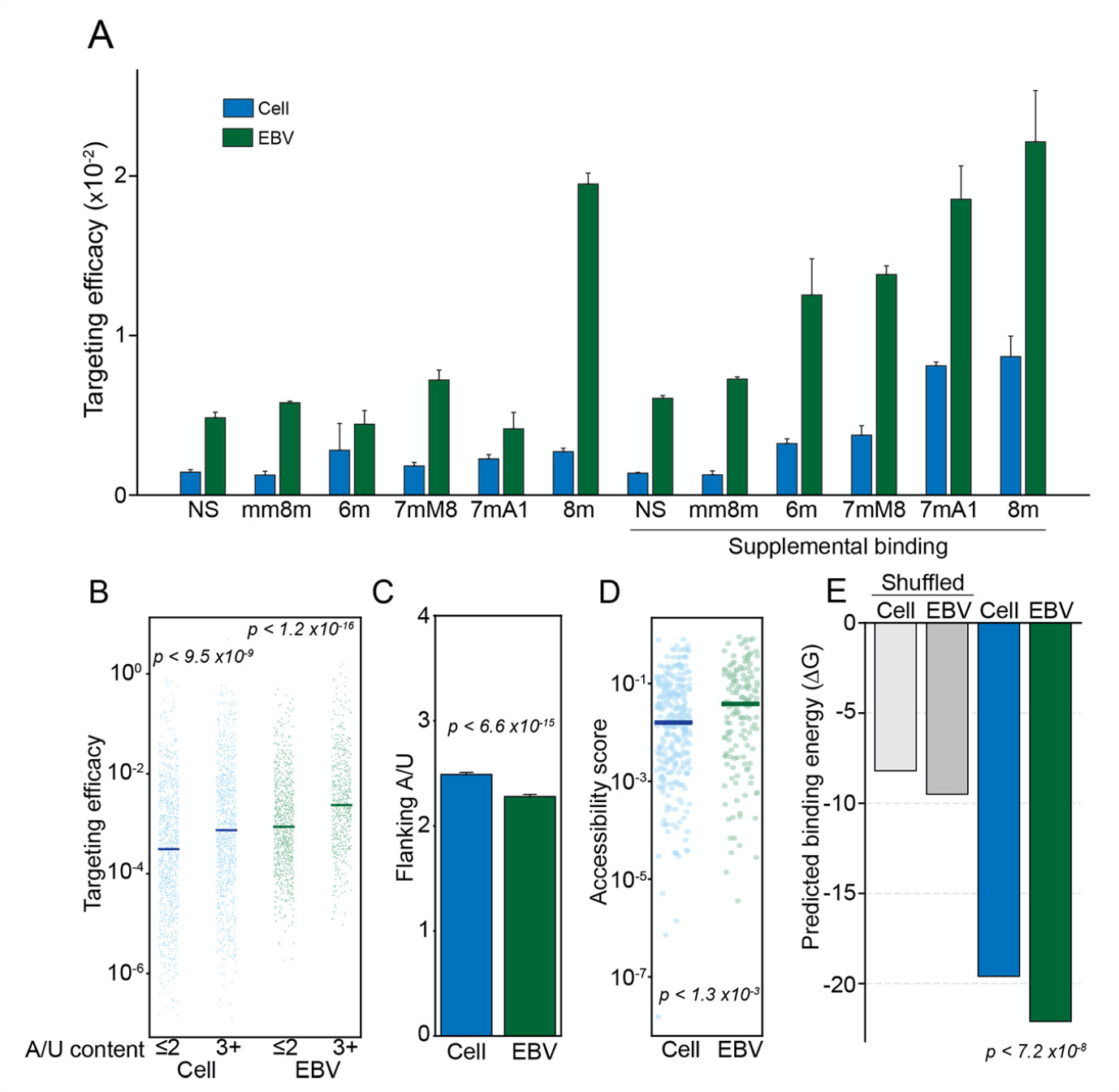
EBV miRNA hybrids are thermodynamically stable. Interactions were categorized by seed match type (NS, no seed; mm8m, mismatch 8mer; 6m, 6mer; 7mM8, 7mer base 8 match; 7mA1, 7mer A1 (A opposite base 1); 8m, 8mer), with “supplemental” interactions requiring >3 bases of complementarity between bases 13-17 of the miRNA and its target. (A) Median targeting efficacy for each type of seed match comparing viral and cellular miRNA interactions. (B) The targeting efficacy of each interaction binned by flanking A/U content of each miRNA target site. The two most proximal bases on each side of the seed binding region were considered. P-values were calculated using the KS test. (C) The mean number of flanking A/Us (max = 4) for cell and EBV miRNA target sites. P value was calculated via KS test. (D) The predicted local site accessibility score of each miRNA target site, using RNAplfold (-W 80 -L 40 -u). The scores indicate the probability that all 14 bases, centered on position 7 of the miRNA target site, will be unpaired. P-values were calculated using the KS test. (E) Predicted minimum free binding energies (ΔG) were calculated for each hybrid using the RNAcofold function of the Vienna RNA Suite(108). As a control, ΔG calculations were performed on shuffled sequences, with 100 permutations performed for each hybrid pair. The ΔG values of cellular and viral hybrids were compared by KS test.

In addition to the extent of seed matching, miRNA binding is improved when target sites are flanked by A/Us(69). Considering the 4 bases directly adjacent to the seed binding site (2 bases on each side of the seed binding region), we found that sites with high A/U content were associated with improved targeting efficacy for both cell and viral miRNAs (*Figure 4B, Figure S10A*). However, EBV, KSHV and MHV68 miRNAs tended to bind sites with fewer flanking A/Us (*Figure 4C, Figure S10B*), indicating that the number of flanking A/Us fails to explain the higher targeting efficacies of viral miRNAs.

Because secondary structure of target sites reduces accessibility and interferes with RISC formation(70), transcript regions with less intramolecular binding propensities are typically more effective miRNA targets. We therefore assessed target site accessibility for each CLASH hybrid using the RNApl-fold(71) subpackage of the Vienna RNA suite using a window size and maximum base pairing separation of 80 and 40 bases, respectively. This analysis revealed that EBV miRNA target sites are, on average, more accessible than the target sites of host miRNAs (*Figure 4D*). This indicates that EBV miRNAs evolved to target more accessible regions of target RNAs providing one likely explanation for the observed higher targeting efficacies of EBV miRNAs.

The last feature that we assessed to determine the molecular basis of enhanced EBV miRNA-mRNA targeting efficacies was predicted minimum free energies of miRNA-mRNA hybrid pairs (across the entire miRNA and its target). This analysis revealed that EBV miRNA-mRNA hybrids tend to form more thermodynamically stable interactions than their cellular counterparts (*Figure 4E*). Extending this analysis to KSHV and MHV68 miRNAs, we found that like EBV miRNAs, targeting efficacies are higher for each seed class (not shown) and we found that KSHV and MHV68 miRNAs generally form more thermodynamically stable interactions than their cellular counterparts (*Figure S10C*). Together, these analyses indicate that *γ*HV miRNAs evolved with nucleotide sequence compositions that favor stronger hybrid interactions, providing another likely explanation for the enhanced targeting efficacy of viral miRNAs.

### EBV miRNAs have a greater impact on their targets than their cellular miRNA counterparts

To test whether the higher overall targeting efficacies of EBV miRNAs translates to increased effectiveness in destabilizing their target mRNAs, we performed correlation analyses between the expression of each viral and cellular miRNA and their respective mRNA targets across EBV positive BL and GC datasets. Correlations were displayed in cumulative distribution plots (*Figure 5*). Shifts in distributions from permuted correlations of the same set of miRNAs with random mRNA targets were analyzed using the Kolmogorov-Smirnov (KS) test (*Figure 5*). In both BLs and GCs, viral miRNAs showed strong inverse correlations with their targets (BL, p < 2.1×10^−6^; GC, p < 1.2×10^−5^). In contrast, correlations between human miRNAs and their targets were less pronounced and not statistically significant. These results provide *in vivo* evidence of functional impacts of EBV miRNAs on targets identified by CLASH and they show that the higher targeting efficacies of EBV miRNAs likely translates into stronger functional influences on their targets.

**Figure 5.**
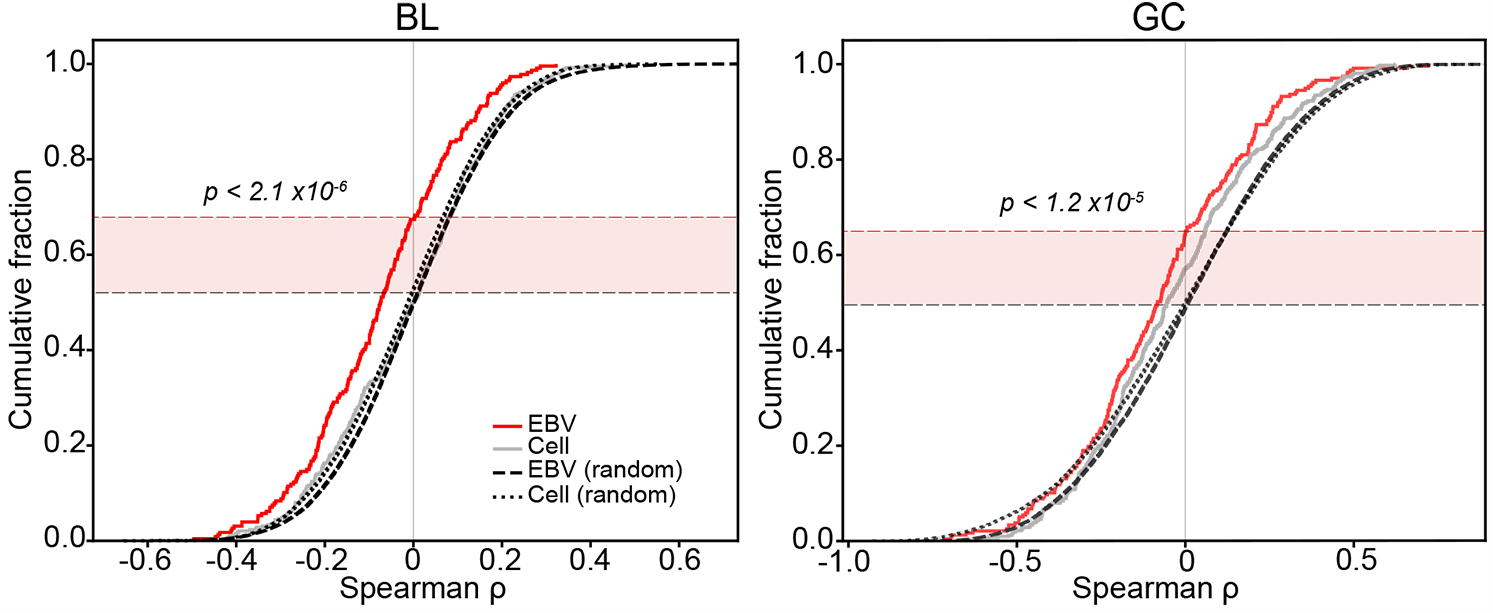
EBV miRNAs and their targets have strong inverse correlations in primary BLs and GCs. For each CLASH miRNA-mRNA hybrid, mRNA and miRNA expression levels (c.p.m.) were correlated across all EBV-positive BL and GC tumors. Correlations were performed on hybrids that met the following criteria: 1. Average miRNA expression in tumors >= 10 c.p.m., 2. Average mRNA expression in tumors > 5 t.p.m., 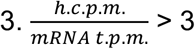, and 4. miRNA-mRNA interactions occurred within the 3’-UTR, where miRNA mediated repression is more often effective(109). Cumulative distribution plots were generated using ranked Spearman correlation coefficients for each species. As a control, expression levels of the miRNA of each hybrid pair analyzed was correlated with expression levels of a random mRNA across the same tumors. For each randomized interaction, 100 permutations were run. Correlation values of EBV miRNA-mRNA pairs were compared to randomized controls; P-values were generated using the KS test.

### EBV miRNA target distributions

In addition to assessing the impact that EBV miRNAs have on their individual targets, we also explored the breadth of viral miRNA targets to investigate the overall impact that EBV miRNAs have on the cell transcriptome. We first quantified the number of targets attributable to each viral miRNA (*Figure 6A*). Several viral miRNAs predominantly target a single transcript (for example, BART3-3p, BART17-3p, and BART20-5p), while others such as BART5-5p, BART7-3p, BART9-3p, and BART16 have many strong targets. The predicted seed-pairing stability (SPS; the predicted binding energy of positions 2-8 of a miRNA bound to its reverse complement target) of miRNAs is a predictor of how promiscuous the miRNA is(72) and these findings are borne out for EBV miRNAs. For example, BART7-3p (AUCAUAG; ΔG° = −5.53 kcal mol^−1^), BART 9-3p (AACACUU; ΔG° = −5.54 kcal mol^-1^), and BART16 (UAGAUAG; ΔG° = −5.73 kcal mol^−1^; *Table S3*) have a relatively low SPS and extensively target a multitude of mRNAs (>100 h.c.p.m.: BART7-3p, 128; BART16, 35; BART9-3p, 25). BART5-5p (AAGGUGA; ΔG° = −7.98 kcal mol^−1^) has an average SPS (ranking in the 48^th^ percentile of all cellular miRNAs) and intermediate, 82 high abundance targets. MiR-NAs with high SPS, such as BART3-3p (GCACCAC; ΔG° = -11.29 kcal mol^−1^), BART17-3p (GUAUGCC; ΔG° = −9.37 kcal mol^−1^), and BART20-5p (AGCAGGC; ΔG° = −11.53 kcal mol^-1^), which have 11, 4, and 1 high abundance target(s), respectively. These high SPS miRNAs tend to extensively target one transcript, with BART3-3p targeting IPO7 over 12-fold more often than its next most frequent interaction partner, BART17-3p:RBM8A accounting for nearly 8-fold more hybrids than its next best target, and BART20-5p:UBE2H representing over 2-fold more hybrids than its next most common target. Overall, despite the relatively focused functions of some EBV miRNAs, EBV miRNAs as a whole tend to display greater breadth in targets than their cellular counterparts (*Figure 6B*). This may be a means through which EBV evolved to influence a larger target set despite having a smaller repertoire of encoded miRNAs.

**Figure 6.**
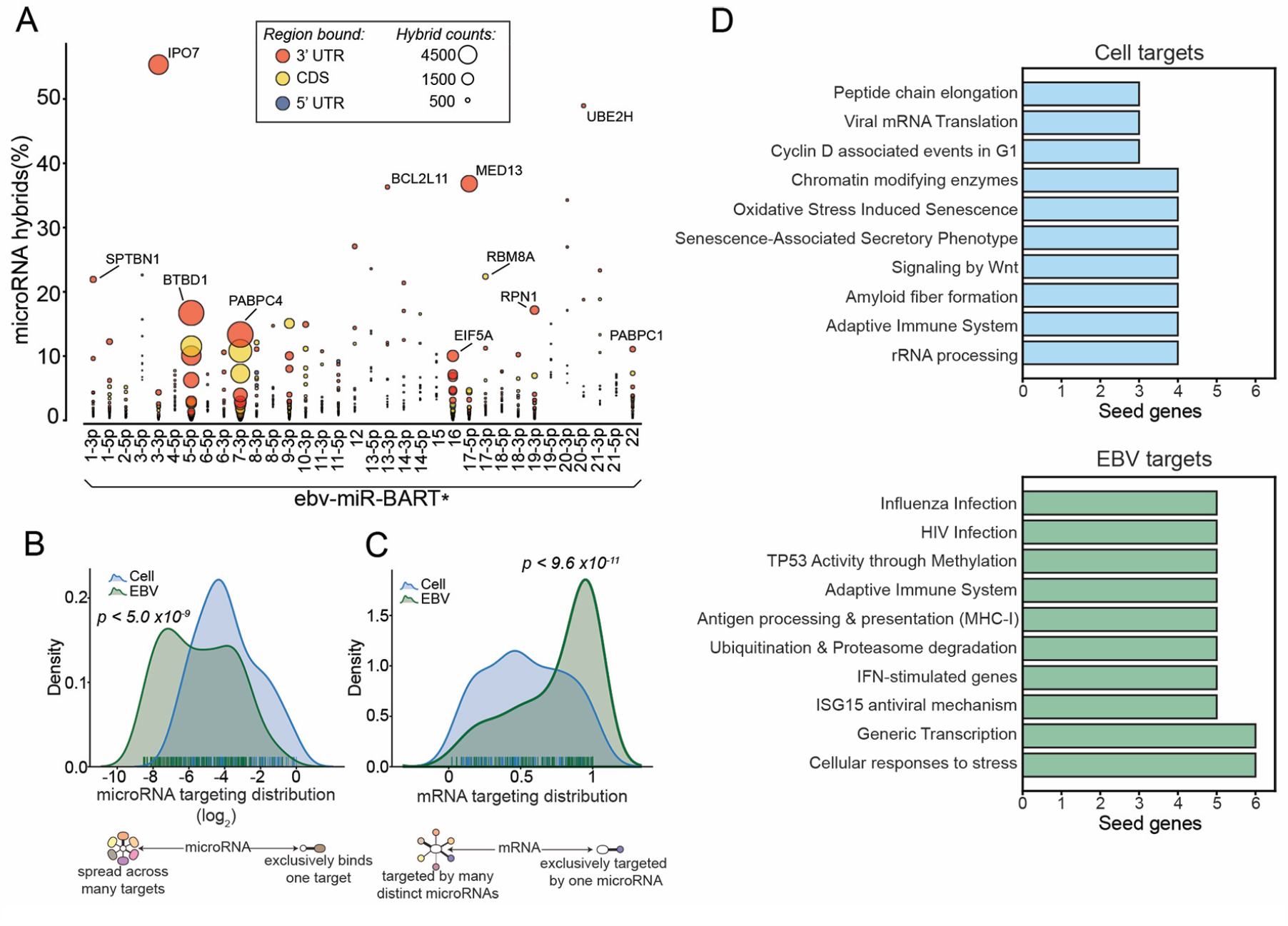
*γ*HV miRNAs target components of the ubiquitin-proteasome system. (A) SNU719 hybrids containing EBV miRNAs. Each interaction was represented by a circle; circle size corresponds to the total number of hybrids formed (*microRNA*_x_:*mRNA_y_ h*.*c*.*p*.*m*); The y-axis values represent the percent of all hybrids that contain the indicated miRNA, 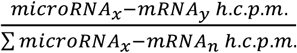. (B) The distribution of y-axis values from (A), extended to all hybrids. Cellular and viral hybrids were compared; P-value was generated using the KS test. (C) The fraction of individual miRNAs hybridizing with each mRNA, 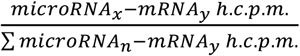., comparing viral and human distributions; P-value was generated using the KS test. (D) Pathways targeted by EBV and cellular microRNAs. Protein-protein interaction networks of each of the top 20 EBV or cellular target genes were obtained from StringDB(52), and resulting protein names were submitted to Enrichr(53) for pathway enrichment analysis (interrogating pathways included in the Reactome database). All statistically significant pathways (FDR < 0.05) were assigned to the target gene.

While EBV miRNA targeting is generally spread across more transcripts, they tend to be directed towards transcripts regulated by fewer miRNAs (*Figure 6C*). This indicates that cellular miRNAs more often work in a cooperative fashion. The higher targeting efficacies of viral miRNAs may reduce their requirement for cooperative targeting, as they tend to target a repertoire of transcripts that are distinct from cellular miRNAs. As an extension of their enhanced targeting efficacy, viral miRNAs are not constrained by the necessity of cooperative targeting, allowing them each to serve a unique purpose, and increasing their influence over host cell transcriptomes.

### EBV miRNAs interfere with immune signaling in the tumor microenvironment

Next, we sought to investigate the potential functional impact of the most influential EBV miRNA-target interactions in modulating the host cell environment. Using the targets of the top 20 EBV miRNA-target interactions as network seed genes, we applied enrichR(53) to query pathway enrichment for each seed gene and its 20 strongest interacting partners. As a control, we performed a similar analysis on the targets of the top 20 cell miRNA-target interactions. This analysis revealed selective enrichment for Influenza and HIV infection, antigen processing and presentation (MHC class I), and IFN-stimulated genes and ISG15 antiviral mechanisms as top EBV miRNA pathway hits (*Figure 6D, Table S4*). This suggests that the EBV miRNA-target interactions within the top 20 highest targeting efficacy interface with adaptive and innate immune response pathways.

Previous studies have shown that EBV positive GCs have higher immune cell infiltration than their EBV negative counterparts(73), likely due to sporadic expression of lytic viral proteins within the tumor. Using CIBERSORTx(74) to infer the immune cell infiltration in each tumor through deconvolution of immune cell signatures within the GC dataset (using only the microsatellite stable (MSS)-only cohort), we found higher levels of T-cell and macrophage subtypes in the EBV positive cohorts (*Figure S11A*). We also determined the diversity of infiltrating T-cells within these tumors by assessing the number of unique T-cell receptor sequences for each tumor. Utilizing MIXCR(75), we realigned each tumor RNA-seq dataset to all potential combinations of rearranged T-cell receptors and used VDJTOOLS(76) for QC and T-cell clone quantifications. This analysis showed that consistent with the finding of higher levels of T-cell infiltration in EBV positive GCs through CIBERSORTx analysis, we found a greater diversity of T-cell clones in EBV positive tumors than in EBV negative tumors (*Figure S11B*). Together, these results confirm the findings of The Cancer Genome Atlas Research Network(56), showing that EBV likely induces some level of adaptive immune response in EBV positive GCs.

While the presence of EBV clearly induces immune cell infiltration into EBV associated tumors, we hypothesized that the interactions of viral miRNAs with immune regulatory pathways in infected cells acts as a counter measure to mitigate immune cell-mediated targeting of virus-infected tumor cells. To functionally assess this possibility we first identified potential direct and indirect targets of viral miRNAs in the EBV+ BL and GC cohorts by correlating the sum of viral miRNA expression values with the expression levels of each cell gene (*Table S5*). Gene Set Enrichment Analysis (GSEA)(77, 78), using the pre-ranked BL and GC miRNA-gene correlations, revealed that viral miRNA expression negatively correlates with the IFN*γ* and TNF*α* signaling pathways (*Figure 7A*). The inverse relationship between IFN*γ* and TNF*α* signaling pathways and EBV miRNA expression suggests that EBV miRNAs mitigate IFN*γ*- and TNF*α*-mediated amplification of the immune response. To more directly assess the relationship between EBV miRNAs and adaptive immune response, we used CIBERSORTx(74) to infer immune cell infiltration in each tumor sample from the BL and GC datasets (*Table S6)*. This analysis showed that CD8 T-cells, CD4 T-cells, and macrophages inversely correlate with viral miRNA expression (*Figure 7B*). Assessing the infiltrating T-cell diversity across these tumors showed a strong inverse correlation between unique T-cell clonotype counts and viral miRNA expression (*Figure 7C, Table S7*), further supporting a role of EBV miRNAs in interfering with the immune response to EBV infection within these tumors. These findings are most tightly associated with EBV miRNAs rather than other expressed EBV transcripts since EBV miRNAs show a stronger inverse relationship with T-cell clonotype numbers than viral pol II transcripts (*Figure S12*). Together these analyses provide evidence that the targeting of components of immune response pathways by EBV miRNAs results in the dampening of innate and adaptive immune responses to viral infection in EBV positive cancers.

**Figure 7.**
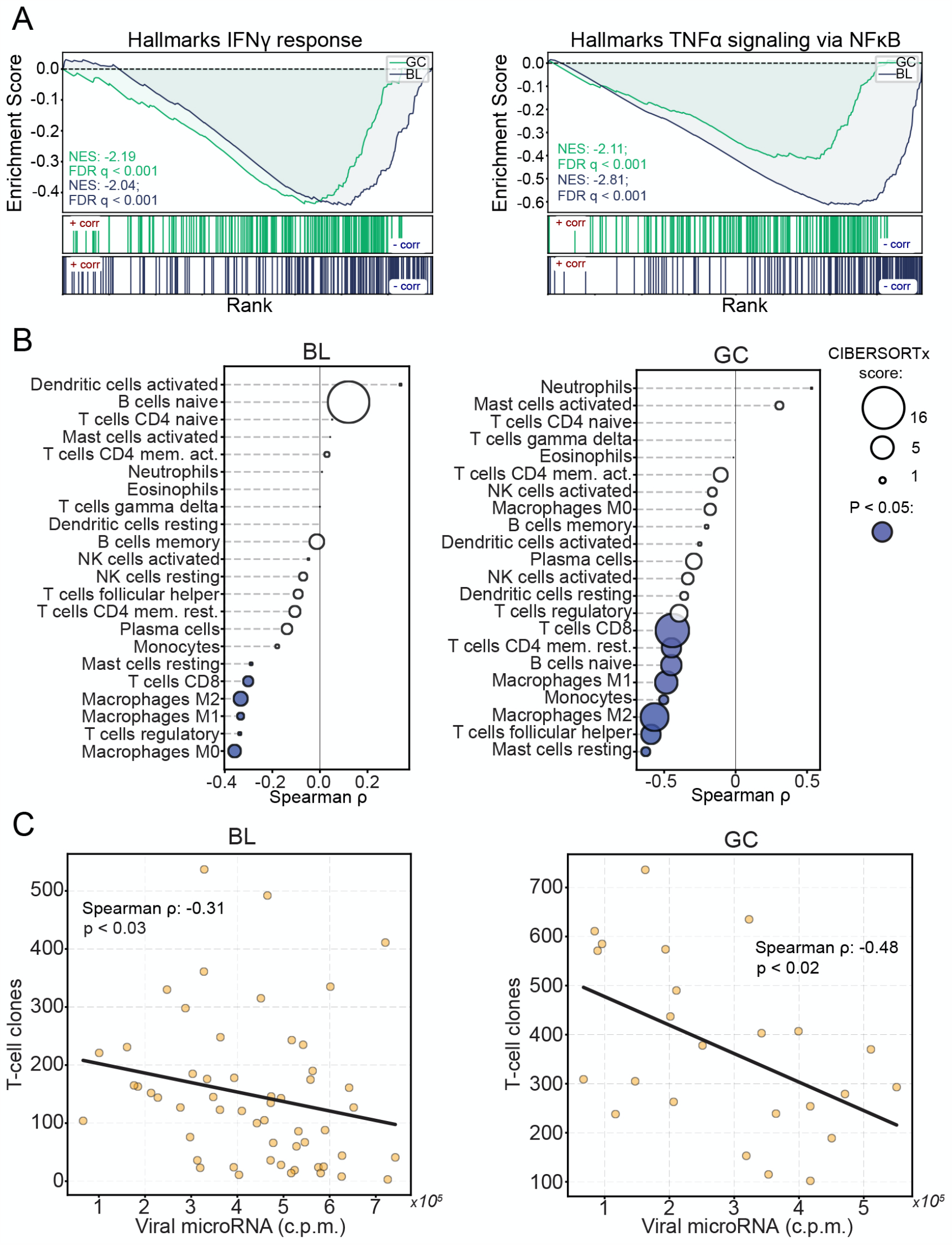
Viral miRNAs interfere with anti-viral immunity. The sum of all viral miRNA c.p.m. was correlated to expression levels of each mRNA (c.p.m.) in EBV-positive BL and GC tumors. The correlations were ranked, then gene lists were interrogated using GSEA. (A) Stacked GSEA curves of the Hallmarks IFN*γ* signaling pathway (left) and the TNF*α*-NFKB signaling pathway (right). NES, nominal enrichment score, FDR, false discovery rate. (B) Immune cell infiltrates were inferred using CIBERSORTx. Each cell type was correlated (spearman) to the sum of EBV miRNAs across EBV-positive BLs and GCs. Circle size represents the average CIBERSORTx absolute score across all tumors, filled circles represent statistically significant (P < 0.05) spearman correlations. (C) T-cell clonotypes were obtained for each EBV-positive tumor sample using MIXCR; The number of unique clonotypes was correlated with the sum of viral miRNAs in EBV-positive BLs and GCs.

## DISCUSSION

By integrating miRNA and mRNA expression levels with hybrid quantifications obtained via CLASH, we used an affinity constant calculation to assess targeting efficacy for each interaction. Comparing targeting efficacies of viral and host miRNA-target interactions, we found that viral miRNAs generally bind their targets more effectively than host miRNAs. The elevated efficacy of EBV miRNA-mRNA interactions transcended seed match and the level of flanking AU content. Instead, we found that the complexes formed between EBV miRNAs and their targets were more thermodynamically stable and tended to occur at target sites with less secondary structure. In metazoans, broadly conserved miRNA target sites tend to facilitate the most effective targeting interactions. However, these sites exhibit a negative selection over time that tempers miRNA binding efficacy. Unlike targets of broadly expressed miRNAs, co-expressed targets of miRNAs that exhibit tissue specific expression tend to evolve effectivity-reducing alterations in target sites(79), suggesting that the major role of miRNAs is to fine-tune gene expression rather than to significantly alter it(80). The continued coevolution of cell miRNAs and their target genes presents an internal arms race that ultimately reduces the impact of the majority of miRNA-mRNA interactions. These constraints do not apply to viral miRNAs which have evolved to effectively target a large number of host transcripts, with some miRNAs such as BART7-3p effectively binding many target mRNAs, and others, such as BART3-3p, homing in on a single transcript (notably, this latter class of viral miRNA might implicate singular targets that are critical to viral persistence and/or expansion in host cells, potentially presenting a vulnerability to the virus). The tendency of viral miRNAs to exhibit high targeting efficacies therefore arm them with the unmatched ability to influence the landscape of host RNA expression.

A longstanding challenge in deciphering miRNA function is the accurate assessment of its targets(81). Because the most favorable interactions require only 7 bases of complementarity, even the most infrequent 3’ UTR binding motifs are widely dispersed throughout the transcriptome. Still, the number of true miRNA binding sites is at least an order of magnitude lower than the number of seed complementary sequences in transcriptome 3’ UTRs. The ability to discern true miRNA targets has improved dramatically with the advent and continued development of binding site prediction algorithms(69, 81), and even more so with high throughput experimental approaches such as PAR-CLIP(82). These tools have seeded studies of EBV miRNA functions, leading to the discovery of consequential mRNA targets(83) including mediators of apoptotic signaling, BBC3 (BART5-5p)(84) and BAD (BART20-5p)(85) and regulators of innate immunity, NDRG1 (multiple)(86), IL12A (BART?), MICB (BART2-5p)(87), IFI30 (BART1 and BART2), LGMN (BART1 and BART2), and CTSB (BART1 and BART2)(88). However, both putative and empirically identified viral miRNA targets have often been difficult to validate(89). In our system, the BBC3 interaction was the 220^th^ most prevalent hybrid of BART5-5p (38 h.c.p.m.), CTSB was the 375^th^ most abundant BART1-5p target (6 h.c.p.m.), and combined, NDRG1 hybrids ranked 3681^st^ among all viral miRNA targets (25 h.c.p.m.). We were unable to detect any viral miRNA hybrids with IFI30, LGMN, or BAD, while MICB and IL12A were not sufficiently expressed in SNU719 cells. Overall, little overlap exists between targets identified among previously published high throughput EBV miRNA targetome studies(89), yet, one consistently identified interaction is that between BART3-3p and IPO7, an interaction that has subsequently been used as a control in studies of EBV miRNA targeting(35, 90), we detect substantial levels of this interaction (6846 h.c.p.m.; ranked 1^st^ among BART3-3p targets), suggesting the quantitative nature of this approach helps distinguish the targets of both viral and host miRNAs that are likely to be impacted the most.

Among the most frequent targets of several EBV miRNAs were transcripts coding for ubiquitin ligases and adapters. These include the E3 ubiquitin ligases, FBXO21 (BART21-5p; ranked 1^st^ among all BART21-5p interactions), TRIM65 (BART7-3p; ranked 5^th^), RNF38 (BART5-5p; ranked 7^th^), TRIM8 (BART16; ranked 1^st^), and KCMF1 (BART1-3p; ranked 2^nd^ and BART3-3p; ranked 2^nd^), the E2 ubiquitin ligase, UBE2H (BART20-5p; ranked 1^st^ and BART7-3p; ranked 17^th^), and the E3 ubiquitin ligase adapters, BTBD1 (BART5-5p; ranked 1^st^) and UBXN7 (BART5-5p; ranked 16^th^). Several of these ubiquitin ligases have important roles in host intrinsic immunity against viral infection. Both FBXO21 and TRIM65 are necessary for the production of type I IFNs in response to viral infection(91, 92). TRIM8 is a potent activator of TNF*α* and NFKB signaling(93), both pathways of which we found to have a strong inverse correlation with viral miRNAs in BLs and GCs (*Figure 7A*).

A more general possible functional consequence of inhibiting ubiquitin ligase expression is interference with MHC class I antigen presentation. Previous studies have shown that EBV miRNAs collectively reduce host cell antigen presentation, culminating in the subversion of CD4+ and CD8+ T-cell surveillance(88, 94, 95). Numerous viruses have evolved various mechanisms to interfere with this pathway. For example, the HIV NEF protein redirects HLA-A and HLA-B transportation(96), the Hepatitis B Virus X protein interferes with proteasomal activity(97), K3 and K5 of KSHV promote MHC-I endocytosis and destruction(98), MHV68 MK3 is a ubiquitin ligase that targets MHC-I for destruction(99), and both the U6 protein of HCMV and EBV encoded BNLF2A inhibit TAP-mediated peptide transport and subsequent MHC loading(100-102). While viral protein expression is restricted in EBV-associated tumors, the presence of the virus still elicits an immune response and mutations that help drive tumors produce neo-epitopes that immune cells may recognize as foreign. The survival of EBV positive cells, including tumors, therefore, requires immune subversion or escape(103). Previous studies have shown that EBV miRNA expression is elevated by inoculation of tissue culture grown EBV positive tumor cells in mice(104). Furthermore, we found that EBV miRNAs inversely correlate with immune cell infiltrates in BLs and GCs, suggesting that an important role of the viral miRNAs is to interfere with immune surveillance, thereby helping facilitate the survival and expansion of the underlying tumor.

Altogether, our findings show that EBV miRNAs are uniquely effective inhibitors of target RNAs through facilitating high *in vivo* expression and the targeting of accessible sites with more favorable miRNA/target binding energies, thereby achieving conserved impacts on immune responses to infected tumor cell populations.

## AVAILABILITY

All source code is available in our GitHub repository (https://github.com/flemingtonlab/ebv_clash).

## ACCESSION NUMBERS

Small fraction and CLASH sequences have been deposited to the NCBI Gene Expression Omnibus (<u>GSE147228</u>). Accession numbers for polyA-selected RNA sequencing reads are <u>GSM1267812</u> (Akata)(105) and <u>GSM1104509</u> (SNU719)(73).

## AUTHOR CONTRIBUTIONS

E.K.F., S.T, R.R., and N.A.U. conceived the project; W.B., and M.K. performed the CLASH experiments, N.A.U., and E.K.F. wrote the scripts, processed and analyzed the data, and wrote and edited the manuscript.

